# The effect of fatty acid binding protein-3 exposure on endothelial transcriptomics

**DOI:** 10.1101/2025.04.16.649114

**Authors:** Hien C. Nguyen, Aman Singh, Christina A. Castellani, Mohammad Qadura, Krishna K. Singh

## Abstract

**Background:** Fatty acid binding protein-3 (FABP3) is released in circulation following myocardial infarction and an increased level of circulatory FABP3 is also reported in peripheral artery disease patients exposing endothelial cells to higher level of FABP3. Recently, loss of endothelial FABP3 was shown to protect endothelial cells against inflammation-induced endothelial dysfunction, however, the effect of FABP3-exposure on endothelial cells is unknown. Accordingly, to study the effect of FABP3-exposure on endothelial cells we performed transcriptomic profiling following recombinant human FABP3 (rhFABP3) treatment to endothelial cells.

**Method:** Cultured human endothelial cells were treated with either vehicle or rhFABP3 (50 ng/ml, 6h) and then RNA-sequencing was performed. Gene expression analysis followed by gene ontology (GO) and Kyoto Encyclopedia of Genes and Genomes (KEGG) pathway analyses were performed to identify differentially expressed genes and affected cellular functions and pathways.

**Results:** Differential gene expression analysis revealed *kinesin family member 26b* (*KIF26B*) as the most upregulated and *survival of motor neuron 2* (*SMN2*) as the most downregulated genes in rhFABP3-treated in comparison to vehicle-treated endothelial cells. Most of the differentially expressed genes were associated with endothelial cell motility, immune response, and angiogenesis. GO and KEGG analyses indicated that rhFABP3 exposure impacts several crucial pathways, prominently “Regulation of leukocyte mediated cytotoxicity” and “Natural killer cell mediated cytotoxicity”, suggesting its involvement in endothelial cell physiology and response mechanisms to cardiovascular stress.

**Conclusion:** This is the first study evaluating rhFABP3-induced transcriptomics in human endothelial cells. Our data reveal novel genes and pathways affected by FABP3-exposure to endothelial cells. Further research is necessary to validate these findings and fully understand FABP3’s role in endothelial biology and in cardiovascular diseases like myocardial infarction and peripheral artery disease.

## Introduction

Lipid-related physiology linked to cardiovascular impacts crucially depends upon the bioavailability of cellular lipids, which is implicated by the role of metabolic syndrome in cardiovascular diseases (CVDs) [1] [2] [3]. Central to the regulation of cellular lipid bioavailability and signalling are a family of intracellular lipid-chaperones, the fatty acid binding proteins (FABPs). Heart-type FABP, or FABP3, mainly known to be expressed in myocardiocytes, is integral to cardiac metabolic homeostasis [1] [2] [4] [5]. Meanwhile, FABP3 can also be found in many other tissues, notably skeletal muscles and, to a lesser extent, the brain, testis, kidneys, adrenal glands, and others [6]. Indeed, the unique function of FABP3 remains complex and unclear [7].

Nonetheless, FABP3 is currently investigated as a potential biomarker for cardiac injuries, having been characterized as follows in both animal models and heart failure patients due to negligible plasma concentration and significantly high cytosolic to plasma ratio at rest; and blood elevation detectable within 30 min of chest pain, peak in a few hours, and returning to baseline via renal clearance, all within 24 h [8] [9] [10]. Peripheral artery disease (PAD) and heart failure are cardiovascular complications of atherosclerosis, a chronic vascular inflammatory disorder characterized by circulatory blockage due to the build-up of lipid-laden plaques in the vascular inner walls, leading to downstream ischemia, hypoxia and organ failures [15]. Atherosclerosis is the primary cause of CVDs and is driven by endothelial dysfunction [16] [17]. Interestingly, a consistent higher level of circulatory FABP3 was reported in PAD patients without any signs of cardiac injury, and FABP3 levels correlated with the severity of PAD [11] [12] [13]. Notably, the same study reported a significant upregulation of FABP3 in skeletal muscle cells in PAD patients compared to healthy individuals. Recently, we reported basal and inflammation-induced expression of FABP3 in endothelial cells; we also demonstrated that endothelial cell-specific loss of FABP3 protects endothelial cells against inflammation-induced endothelial dysfunction and apoptosis [14]. Overall, these findings suggest that FABP3 releases are not exclusive to cardiac injury and may signal earlier cardiovascular events.

Single layer of endothelial cells comprises the inner luminal walls of virtually all blood vessels. Endothelial cells are versatile, in direct contact with blood, and establish a delicate semi-permeable blood-tissue barrier known to extensively regulate selective exchanges and vascular homeostasis at varying capacities across organ systems. They oversee the production of signalling agents that maintain or mediate vasotone (vasodilation vs. vasoconstriction), vessel compliance, barrier/exchange permeability, blood fluidity, inflammation, wound healing, angiogenesis, and thrombosis [18]. In CVD pathogenesis, endothelial cells are often impaired by various stresses, and stressed endothelial cells are activated into a hyper-functional state to alleviate the source of stress. Prolonged endothelial activation causes endothelial dysfunction, featuring a leaky and oxidative barrier that exacerbates injuries and propagates the damaging agents, a hyper-inflammatory environment leading to chronic inflammation, dysregulated metabolism, diminished vasotone, and impaired vascular homeostasis [19].

Given the remarkable capacity of fatty acid-binding protein 3 (FABP3) as a biomarker, albeit not specific to cardiac injury, and that endothelial cell are one of the first cells to be exposed to elevated levels of circulatory FABP3 in conditions like heart failure and PAD, the impacts of circulatory FABP3 on endothelial cells and endothelial function are yet to be elucidated. As endothelial dysfunction is central in atherosclerosis and CVD, the mechanisms by which FABP3 influences endothelial function warrant an investigation. This study explores the transcriptomic profiles of endothelial cells subjected to FABP3 exposure under the hypothesis that circulatory FABP3 regulates endothelial function. We aim to pursue insights into the link between FABP3 and endothelial dysfunction, potentiating the development of novel clinical applications of FABP3 in the cardiovascular field.

## Materials and Methods

### Cell Culture and Treatment

Human umbilical vein endothelial cells (HUVECs, pooled, Lonza; passage 4) were cultured in endothelial cell growth medium-2 (EGM-2 Bulletkit; Lonza) supplemented with growth factors, serum, and antibiotics at 37^0^C in humidified 5% CO2. Confluent HUVECs were maintained in six-well plates, starved overnight before treating with either a vehicle (PBS) or rhFABP3 (50 ng/ml; Cayman Chemical) for 6 hours.

RNA sequencing (RNA-seq) and Analysis **-** Total RNA was extracted from HUVECs using TRIzol (Invitrogen) reagent and quantified with the NanoDrop ND-1000 spectrophotometer as described. RNA quantity and purity were assessed with the NanoDrop ND-1000. RNA sequencing was performed at The Centre for Applied Genomics, The Hospital for Sick Children, Toronto, Canada. Sequencing was conducted on the Illumina HiSeq2500 platform, using the NEBNext Ultra II Directional RNA Library Prep Kit for Illumina (E7760; New England Biolabs) and bcl2fastq2 v2.20 for paired-end reads (125 base pairs). Reads were generated in FASTQ format and, on the Compute Canada platform, subjected to 1) trimming of low-quality reads using Trimmomatic based on the adapters TruSeq3-PE [20], 2) quality assessment using FASTQC^1^, and 3) Kallisto transcriptome pseudoalignment using the GRCh38 (Genome Reference Consortium human genome build 38) indices [21] [22] [23]. Low-quality counts in RNA sequencing data are identified and filtered out during the preprocessing steps to ensure data accuracy before differential expression analysis. Raw reads are subjected to Trimmomatic to trim low-quality sections, such as the adapter sequences TruSeq3-PE. Reads with a high proportion of ambiguous base and those trimmed to a length below thresholds are considered low quality and are discarded. The quality of the trimmed reads is also assessed using FASTQC, which generates reports on base quality, GC content, and the presence of sequencing artifacts. Differential gene expression analysis was conducted using edgeR on R version 4.3.1 [24] [25]. P-values were generated using edgeR’s model for discrete count data that includes dispersion estimation, and the Benjamini-Hochberg method was applied for false discovery rate (FDR) adjustment. Gene Ontology (GO) and Kyoto Encyclopedia of Genes and Genomes (KEGG) pathway analyses were performed using functions goana and kegga in the limma package [26]. To enable a broader assessment of biological mechanisms given a relatively low number of differentially expressed genes following FDR correction, genes with raw p-values < 0.05 and log (2) fold change of lesser or greater than -1 or 1 from edgeR were considered for GO and KEGG analysis, acknowledging the trade-off of potentially increasing the rate of Type-1 errors [27] [28].

## Results

Quality Assessment **-** RNA integrity, quantity, and purity were assessed with the NanoDrop ND-1000. The A260 /A280 optical density (OD) ratios yielded values of about 2.0, which confirmed the purity of our RNAs (S1 Table). The intensity of the 28S ribosomal RNA was about twice (indicated by % of total Area) that of the 18S ribosomal RNA, confirming the integrity of RNA (S1 and S2 Figs) used in this study. These results indicate that the RNA used to perform RNA-seq were not degraded and pure.

FABP3 Exposure Induced Differential Gene Expression in Endothelial Cells **-** The impact of rhFABP3 exposure on HUVECs’ gene expression was examined via *RNA-seq*. Genome-wide differential gene expression (DGE) of 15,688 genes in rhFABP3-treated HUVECs vs. vehicle control cells were assessed following removal of low-quality counts. Principal component analysis (PCA) on the first and second principal components, capturing 58.71% and 20.7% variance, respectively, showed distinct samples clustering between the rhFABP3-treated and vehicle-treated groups in PC1, suggesting changes in global gene expression profiles in HUVECs due to rhFABP3 exposure (Fig 1A). DGE analysis results visualized using Volcano and Manhattan plots identified 11 genes with significant differential expression that satisfy a log (2) fold-change threshold of 1 and -1 (equivalent to one doubling of gene expression up or downward) in rhFABP3-treated group vs. vehicle controls; genes with FDR-adjusted p-values < 0.05 are considered significantly differentially expressed, and of these genes, 7 genes were upregulated and 4 were downregulated (Figs 1B and 1C) (Table 1). Accordingly, differentially expressed genes are distributed between chromosomes 1, 2, 5, 6, 12, 16, 19 and 21, and the *IMPDH1P10* processed-pseudogene is the most upregulated (log2 fold-change of 8.75), followed by the protein-coding genes: *KIF26B* (7.10), *NCR1* (4.17), and *DNAJC14* (1.95). *CENPBD1P* pseudogene is the most downregulated (log2 fold-change of -8.98), followed by the protein-coding genes: *ENSG00000269242* (-3.45), *CFAP298-TCP10L* (-3.10), and *SMN2* (-2.28).

**Table 1:** Summary of FDR-significant (FDR < 0.05) top-differentially-expressed genes by at least log (2) fold-change of 1, including top up- and down-regulated DE protein-coding genes, in HUVECs treated with rhFAB3 (50 ng/ml, 6h) vs. vehicle controls. Human Genome Organization (HUGO) classification utilized in the gene annotation database. A total of 15688 genome-wide genes were validated from processed RNA-seq results and tested for differential gene expression. Gene names are derived from genecards.org. FDR-significant (p < 0.05) top-differentially-expressed genes by at least log (2) fold-change of 1.

| *Top up-regulated differentially-expressed genes* |  |  |  |
| --- | --- | --- | --- |
| HUGO Gene | Gene name | Fold Change | p-value |
| KIF26B | *Kinesin Family Member 26B* | 137.08 | 7.16E-07 |
| NCR1 | *Natural Cytotoxicity Triggering Receptor 1* | 18.02 | 3.89E-05 |
| DNAJC14 | *DnaJ Heat Shock Protein Family (Hsp40) Member* | 3.86 | 4.31E-02 |
| CCDC125 | *Coiled-Coil Domain Containing 125* | 2.68 | 2.90E-02 |
| IMPDH1P10 | *Inosine Monophosphate Dehydrogenase 1 Pseudogene 10* | 431.03 | 2.87E-10 |
| MICE | *Unnamed* | 146.26 | 9.12E-07 |
| PKD1P3 | *Polycystin 1, Transient Receptor Potential Channel Interacting Pseudogene 3* | 3.07 | 5.55E-03 |
| SMN2 | *Survival Of Motor Neuron 2, Centromeric* | 0.21 | 8.64E-03 |
| *Top down-regulated differentially-expressed genes* |  |  |  |
| CFAP298-TCP10L | *CFAP298-TCP10L Readthrough* | 0.12 | 2.02E-03 |
| Unnamed | *Glycosyl Hydrolases Family 38 C-Terminal Beta Sandwich Domain-Containing Protein* | 0.09 | 3.44E-05 |
| CENPBD1P | *CENPB DNA-Binding Domain Containing 1, Pseudogene* | 0 | 1.02E-07 |

**Fig 1.** Differential gene expression (DGE) analysis of HUVECs treated with rhFABP3 (50ng/ml) for 6 hours vs. Vehicle. **(A)** Principal Component Analysis (PCA) plot clustering the samples of HUVECs treated with rhFABP3 (50ng/ml) for 6 hours vs. Vehicle (red = rhFABP3; blue = Vehicle), accessing their global expression of 15688 genes derived from RNA-sequencing results; the x- and y-axes represent the first and second principal components, which capture the most (58.71%) and second-most (20.70%) variance within the data, respectively. Volcano (B) and Manhattan (C) plots of DGE genes in HUVECs treated with rhFABP3 (50ng/ml) for 6 hours vs. vehicle controls. **(B)** Log (2) fold change is plotted against –log (10) false discovery rate (FDR) adjusted p-values; genes with FDR-adjusted p-values less than 0.05 (dashed y-intercept) that pass the log (2) fold-change of 1 or -1 (dashed x-intercepts) in differential expression are labelled (red). **(C)** DGE genes tested are localized to their chromosomes (x-axis); genes with FDR-adjusted p-values less than 0.05 (dashed y-intercept) are highlighted (red). N = 3. HUVEC = Human Umbilical Vein Endothelial Cells; rhFABP3 = recombinant human FABP3, MT = mitochondria.

FABP3 Exposure Affects Potential Biological Functions/Pathways in Endothelial Cells **-** Gene ontology (GO) and Kyoto Encyclopedia of Genes and Genomes (KEGG) pathway analyses were conducted to assess the biological significance of the observed differential gene expression in rhFABP3-treated HUVECs. Due to the low number of significant genes meeting our FDR/fold-change filter, we performed GO/KEGG analysis with 80 genes (38 upregulated and 42 downregulated) selected using the unadjusted edgeR p-values < 0.05 and log (2) fold change of lesser or greater than -1 or 1, respectively, against a total of 15,688 tested genes. The analyses revealed several cellular functions and pathways potentially impacted by rhFABP3 exposure, suggesting a multifaceted role of FABP3 in endothelial cell physiology (Tables 2 and 3). GO and KEGG results are predominantly related to immune response and cell cytotoxicity biological processes for the upregulated differentially expressed genes, with the most significant being “Regulation of leukocyte mediated cytotoxicity” and “Natural killer cell mediated cytotoxicity”, respectively. On the other hand, the downregulated DE genes are associated with GO and KEGG terms of complex regulatory implication, including “RNA processing and inflammatory and immune systems mechanisms, with SMN complex (cellular component)” and “NOD-like receptor signaling” pathway being the most significant. Further, a broader assessment of gene expression patterns is illustrated on a heatmap generated from 231 protein-coding genes with edgeR-derived unadjusted p-values < 0.05, showing distinct clusters of differential expression patterns in rhFABP3-treated HUVECs (S2 Table, S3 Fig). These findings illustrate the specific gene expression signatures associated with FABP3 exposure and underscore the potential functional impacts on endothelial cells.

**Table 2:**
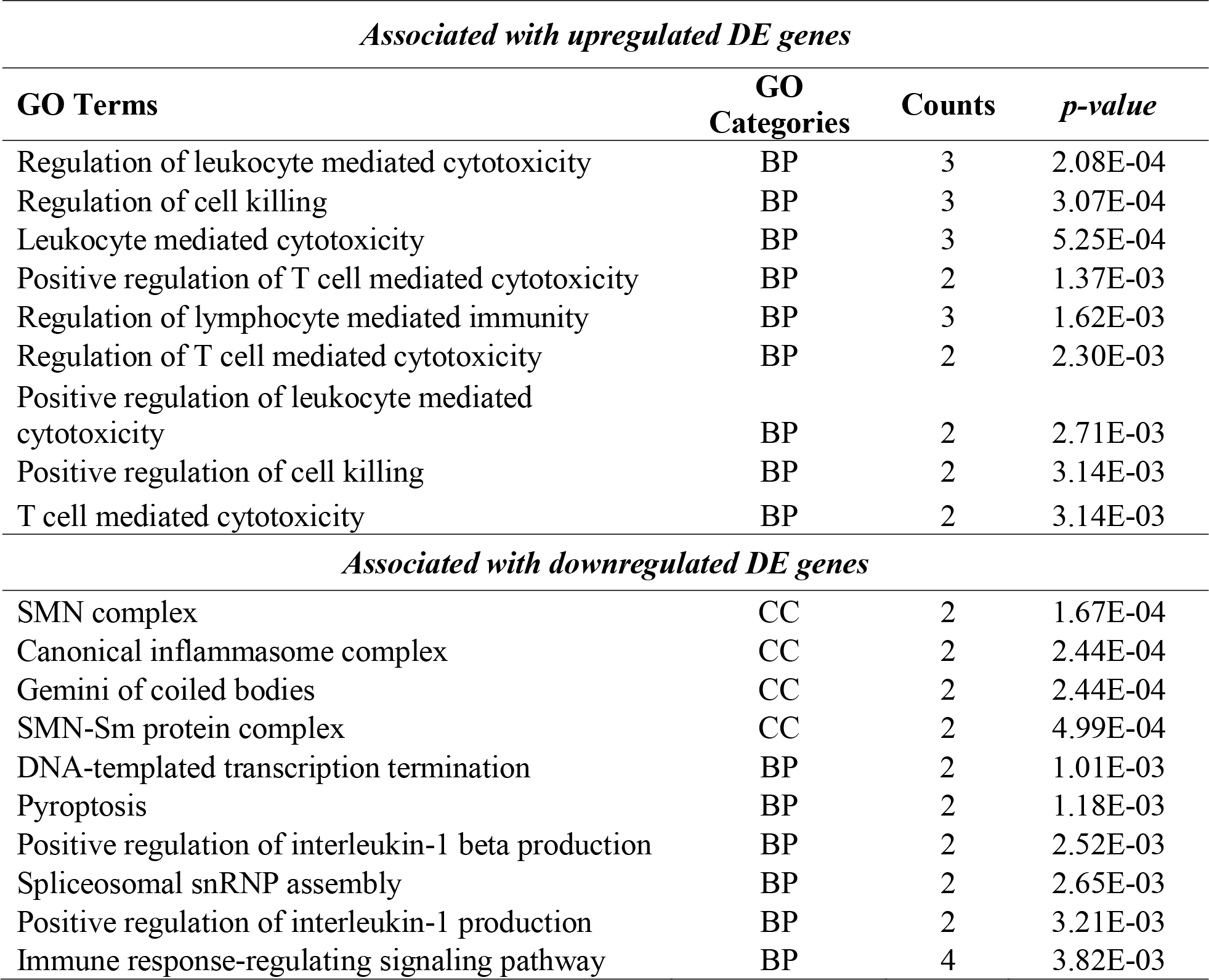
Top up- and down-regulated gene ontologies impacted in HUVECs under rhFABP3 exposure. BP = Biological processes; CC = Cellular Component. Gene ontology (GO) analysis is conducted by enriching 38 genes with unadjusted p-values less than 0.05 and a *log2* fold-change of at least 1 or -1 (one gene expression doubling increase or decrease) against 15688 RNA-seq genes tested for differential gene expression.

| *Associated with upregulated DE genes* |  |  |  |
| --- | --- | --- | --- |
| GO Terms | GO Categories | Counts | *p-value* |
| Regulation of leukocyte mediated cytotoxicity | BP | 3 | 2.08E-04 |
| Regulation of cell killing | BP | 3 | 3.07E-04 |
| Leukocyte mediated cytotoxicity | BP | 3 | 5.25E-04 |
| Positive regulation of T cell mediated cytotoxicity | BP | 2 | 1.37E-03 |
| Regulation of lymphocyte mediated immunity | BP | 3 | 1.62E-03 |
| Regulation of T cell mediated cytotoxicity | BP | 2 | 2.30E-03 |
| Positive regulation of leukocyte mediated cytotoxicity | BP | 2 | 2.71E-03 |
| Positive regulation of cell killing | BP | 2 | 3.14E-03 |
| T cell mediated cytotoxicity | BP | 2 | 3.14E-03 |
| *Associated with downregulated DE genes* |  |  |  |
| SMN complex | CC | 2 | 1.67E-04 |
| Canonical inflammasome complex | CC | 2 | 2.44E-04 |
| Gemini of coiled bodies | CC | 2 | 2.44E-04 |
| SMN-Sm protein complex | CC | 2 | 4.99E-04 |
| DNA-templated transcription termination | BP | 2 | 1.01E-03 |
| Pyroptosis | BP | 2 | 1.18E-03 |
| Positive regulation of interleukin-1 beta production | BP | 2 | 2.52E-03 |
| Spliceosomal snRNP assembly | BP | 2 | 2.65E-03 |
| Positive regulation of interleukin-1 production | BP | 2 | 3.21E-03 |
| Immune response-regulating signaling pathway | BP | 4 | 3.82E-03 |

**Table 3:** Top up- and down-regulated functional pathways impacted in HUVECs under rhFABP3 exposure. Kyoto Encyclopedia of Genes and Genomes (KEGG) based pathways analysis is conducted by enriching 42 genes with unadjusted p-values less than 0.05 and a *log2* fold-change of at least 1 or -1 (one gene expression doubling increase or decrease) against 15688 RNA-seq genes tested for differential gene expression.

| *Upregulated Pathways* |  |  |
| --- | --- | --- |
| **KEGG Pathway Terms** | **Counts** | ***p-value*** |
| Natural killer cell mediated cytotoxicity | 2 | 9.17E-03 |
| JAK-STAT signaling pathway | 2 | 1.63E-02 |
| *Downregulated Pathways* |  |  |
| NOD-like receptor signaling pathway | 2 | 2.92E-02 |

## Discussion

Our RNA-seq DGE analyses identify several significantly impacted genes in rhFABP3-treated HUVECs, which underline the complex nature of endothelial cells response to circulatory FABP3. Of the upregulated genes, *KIF26B* is an oncogene that has been studied in breast, gastric, colorectal, and hepatocellular cancers; its upregulation correlates with risk of metastases, stage progression, and poor prognosis, suggesting capacities as a biomarker [29]. KIF26B is regulated by miR-372 [30] and essential in developmental processes, implicated in the adhesion and polarization of mesenchymal cells [31]. In cancer, KIF26B is involved in the VEGF signaling pathway that prompts angiogenesis [32]. The upregulation of KIF26B in rhFABP3-treated endothelial cells suggests induced mobilization of cellular motility and possibly angiogenesis, suggesting a regulatory metabolic impact of FABP3 exposure on endothelial cells. It is also important to note that FABP3 is released in circulation mainly during ischemic/hypoxic stress [2] and increased angiogenesis may be a compensatory response of endothelial cells. Upregulated *NCR1* gene encodes an activating receptor on natural killer cells, which imposes innate cytotoxicity and surveillance against bacteria, virally infected cells, and tumor cells [33]. NCR1 is known to mediate the pathogenesis of cancer, autoimmune disorders, and infectious diseases, being a target for immunomodulation and immunotherapy [33]. Regulatory factors of NCR1 include cytokines, transcription factors, microRNAs, and post-translational modifications [34] [35] [36] [37]^2-5^. While the gene’s expression is a main feature of natural killer cells, NCR1 has been found in other cell types, such as T-cells [38]. Endothelial NCR1 is poorly understood, and our detection of NCR1’s expression in endothelial cells suggests a novel regulatory link between the innate immune system and the endothelium. Particularly, NCR1 upregulation in HUVECs indicates stimulation of the innate immune system by FABP3 exposure. Next, upregulated gene *DNAJC14* encodes a member of the DNAJ family of intracellular heat-shock chaperone proteins, which are engaged in the cellular stress response and protein quality controls [39]. In particular, they interact with the Hsp70 chaperone proteins via the distinguishing J-domain and assist Hsp70 in re-folding misfolded proteins [40]. While the specific roles of DNAJC14 remain under investigation, aberrant expression of DNAJC14 has been implicated in multiple diseases, including viral infections and neurodegenerative diseases, particularly in the context of misfolded proteins [41] [42]. The upregulation of DNAJC14 in endothelial cells under FABP3 exposure indicates stress response mechanisms, reinforcing the metabolic impact of circulatory FABP3 on the endothelium. Among the downregulated genes *SMN2*, canonically crucial in motor neuron functions and spinal muscular atrophy, which encodes a more truncated and less functional protein than the full-length version expressed by SMN1 [43]. SMN proteins mediate the assembly of small nuclear ribonucleoproteins of spliceosomes, thereby regulating RNA splicing, post-transcriptional processing, and non-coding RNAs [44]. SMN2 downregulation in rhFABP3-treated endothelial cells indicates a negative modulation of RNAs in the endothelium, suggesting a regulatory effect on transcript levels. The rest of the identified DE genes are less well characterized. CCDC125 encodes for a protein that is not well-characterized and may be involved in cellular motility according to its Uniprot profile. The human genome GRCh37 ensembl profile of CFAP298-TCP10L (ENSG00000265590) indicates that it is a protein-coding readthrough transcription between the neighboring chromosome 21 open reading frame 59 and TCP10L (t-complex 10 like) that hasn’t been investigated for any functions. Glycosyl Hydrolases Family 38 C-Terminal Beta Sandwich Domain-Containing Protein (ENSG00000269242) is a novel transcript with gene ontology annotations related to carbohydrate binding and mannosidase activities, according to its gene-card profile. IMPDH1P10, ENSG00000251581, PKD1P3, and CENPBD1P are pseudogenes that remain functionally elusive. Overall, although some DE genes suggest a metabolic and immunity-based response in rhFABP3-treated endothelial cells, a notable amount are pseudogenes, and more than half are currently not characterized. Future validation, such as via qPCR, and characterization of the identified DE genes are necessary to establish more robust mechanistic implications.

A total of 11 significantly differentially expressed genes were identified from our RNA-seq and differential gene analyses (a total of 15688 genes tested) that meet a log (2) fold-change of 1.0 cut-off. This presents an ostensive limitation which may be explained by the small sample size of the study and stringent cutoffs employed. The limitation may be attributed to our low dose of FABP3 (50 ng/ml for 6h). In a study that evaluated 2287 patients with acute coronary syndromes, 332 patients (14.5%) were found with elevated circulatory H-FABP levels (>8 ng/mL). This elevation was associated with an increased risk of death and major cardiac events through a 10-month follow-up period, including recurrent myocardial infarction and congestive heart failure. From the elevated H-FABP cohort, the median level of H-FABP3 in circulation was 16 ng/mL, ranging from 8 to 434 ng/mL [45]. In our previous PAD patients’ study that identified a robust positive correlation between the severity of PAD and blood FABP3 levels, severe PAD patients (ABI < 0.4) exhibited up to an average of 7.22 ng/ml of blood FABP3 [11]. Therefore, our 50 ng/ml of FABP3 is informed by existing clinical data, fitting within these variable ranges of circulatory FABP3 reported in human patients. However, from our study on the loss of FABP3 in endothelial dysfunction, 200 ng/ml of rhFABP3, but not 50 ng/ml, was found to exacerbate ICAM1 and VCAM1 upregulation in HUVECs stressed by LPS for 6h [14]. This not only suggests a negative inflammatory role of FABP3 exposure but also that our current FABP3 dose may fall short in inducing a pronounced gene expression response, overall implying that a higher dose should be considered within the provided clinical ranges for future investigation. Nevertheless, our suggestive previous findings substantiated the current attempt to analyze the total RNAs from rhFABP3-treated endothelial cells using RNA-seq. Hence, the transcriptomic analysis was conducted at our selected dosing regimen to clarify how endothelial cells are affected by FABP3 exposure. We used human umbilical vein endothelial cells in this study, which is the most characterized and an established model to study endothelial biology in vitro [46]. However, it is important to note that endothelial cells are differentiated differently depending on their location [47] [48] and, accordingly, future studies should be performed in endothelial cells of aortic, micro- and macro-vasculature origin.

## Conclusion

In conclusion, the endothelial genome-wide alterations in response to FABP3 exposure were delineated. Differentially gene analyses highlighted several genes associated with endothelial cells metabolic and immune-related stress response invoked by FABP3 treatment, suggesting that FABP3 releases during cardiovascular events impact endothelial function. While the roles of less characterized genes remain to be elucidated, we provide a transcriptomic profile for future research into FABP3’s impact on endothelial biology. Our analysis’s sensitivity and specificity underscore the need for validation in larger sample sizes and with additional experimental treatments. Given that endothelial cells form the first interaction with circulatory FABP3, future studies should expand on these findings, exploring the therapeutic and diagnostic applications of FABP3 within the vascular system.

## Acknowledgements

Conceptualization, K.K.S., and M.Q.; carried out the experiments, H.C.N. and A.S.; data analysis, H.C.N. and C.A.C.; resources, K.K.S.; writing original draft, H.C.N., and K.K.S.; review and editing, H.C.N., A.S., C.A.C., and K.K.S.; supervision, K.K.S.; and funding acquisition, K.K.S. All authors have read and agreed to the published version of the manuscript.

