## Supplementary Files for "The effect of fatty acid binding protein-3 exposure on endothelial transcriptomics"

#### Supplementary Information

##### Supplementary Figure 1 (S1 Fig)

Assay Class: Eukaryote Total RNA Nano
Data Path: Z:\...Eukaryote Total RNA Nano_DE04103349_2023-10-19_13-13-47.xad
Created: 10/19/2023 1:13:47 PM
Modified: 10/19/2023 2:54:02 PM

**Electropherogram Summary**

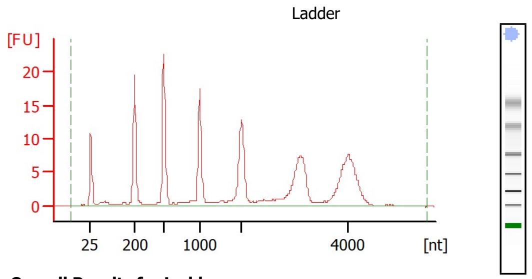


**Overall Results for Ladder**

| | |
|---|---|
| RNA Area: | 214.7 |
| RNA Concentration: | 150 ng/µl |
| Result Flagging Color: | |
| Result Flagging Label: | All Other Samples |

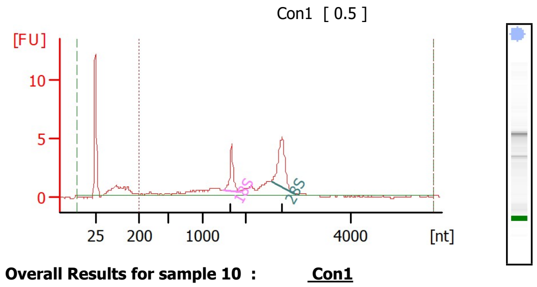


**Overall Results for sample 10 : Con1**

| | |
|---|---|
| RNA Area: | 46.2 |
| RNA Concentration: | 32 ng/µl |
| rRNA Ratio [28s / 18s]: | 1.8 |
| RNA Integrity Number (RIN): | 7.8 (B.02.11) |
| Result Flagging Color: | |
| Result Flagging Label: | RIN: 7.80 |
| Corr. Area 1: | 37.9 |

**Fragment table for sample 10 : Con1**

| Name | Start Size [nt] | End Size [nt] | Area | % of total Area |
|---|---|---|---|---|
| 18S | 1,497 | 1,842 | 5.3 | 11.5 |
| 28S | 2,494 | 2,915 | 9.8 | 21.1 |

**Region table for sample 10 : Con1**

| Name | From [nt] | To [nt] | Corr. Area | % of Total | Color |
|---|---|---|---|---|---|
| DV200 | 200 | 5,552 | 37.9 | 82 | |

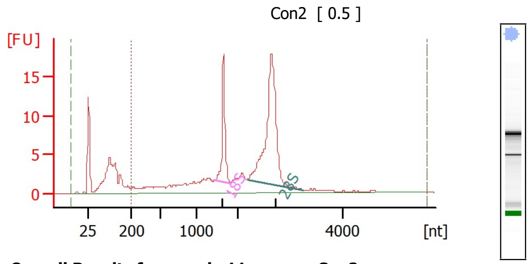


**Overall Results for sample 11 : Con2**

| | |
|---|---|
| RNA Area: | 188.7 |
| RNA Concentration: | 132 ng/µl |
| rRNA Ratio [28s / 18s]: | 2.9 |
| RNA Integrity Number (RIN): | 8.5 (B.02.11) |
| Result Flagging Color: | |
| Result Flagging Label: | RIN: 8.50 |
| Corr. Area 1: | 147.8 |

**Fragment table for sample 11 : Con2**

| Name | Start Size [nt] | End Size [nt] | Area | % of total Area |
|---|---|---|---|---|
| 18S | 1,442 | 1,832 | 18.7 | 9.9 |
| 28S | 2,216 | 3,233 | 54.9 | 29.1 |

**Region table for sample 11 : Con2**

| Name | From [nt] | To [nt] | Corr. Area | % of Total | Color |
|---|---|---|---|---|---|
| DV200 | 200 | 5,595 | 147.8 | 78 | |

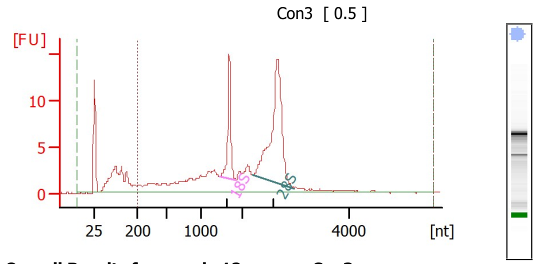


**Overall Results for sample 12 : Con3**

| | |
|---|---|
| RNA Area: | 174.1 |
| RNA Concentration: | 122 ng/µl |
| rRNA Ratio [28s / 18s]: | 3.0 |
| RNA Integrity Number (RIN): | 8 (B.02.11) |
| Result Flagging Color: | |
| Result Flagging Label: | RIN:8 |
| Corr. Area 1: | 144.2 |

**Fragment table for sample 12 : Con3**

| Name | Start Size [nt] | End Size [nt] | Area | % of total Area |
|---|---|---|---|---|
| 18S | 1,432 | 1,822 | 15.4 | 8.9 |
| 28S | 2,195 | 2,968 | 47.0 | 27.0 |

**Region table for sample 12 : Con3**

| Name | From [nt] | To [nt] | Corr. Area | % of Total | Color |
|---|---|---|---|---|---|
| DV200 | 200 | 5,616 | 144.2 | 83 | |

**S1 Fig. Confirmation of RNA quantity and purity for the control-treatment samples (Con1, Con2, and Con3).** Sample Con1 has an RNA integrity number (RIN) of 7.8 and a 28S/18S ratio of 1.8. Sample Con2 exhibits a RIN of 8.5 and a 28S/18S ratio of 2.9. Sample Con3 has a RIN of 8.0 and a 28S/18S ratio of 3.0. Standardized RIN values range from 1 (completely degraded) to 10 (intact, high-quality RNA), and a RIN > 7 is considered suitable for downstream applications. Standardized 28S/18S ratio, which is the relative abundance of the two ribosomal RNA subunits of around 2.0 indicates qualified RNA purity and integrity.

### Supplementary Figure 2 (S2 Fig)

| | | | |
|---|---|---|---|
| Assay Class: | Eukaryote Total RNA Nano | Created: | 10/19/2023 2:51:30 PM |
| Data Path: | Z:\...Eukaryote Total RNA Nano_DE04103349_2023-10-19_14-51-30.xad | Modified: | 10/19/2023 3:14:35 PM |

**Electropherogram Summary**

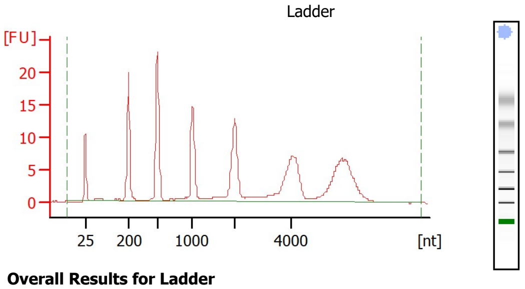


**Overall Results for Ladder**

| | |
|---|---|
| RNA Area: | 221.3 |
| RNA Concentration: | 150 ng/µl |
| Result Flagging Color: | |
| Result Flagging Label: | All Other Samples |

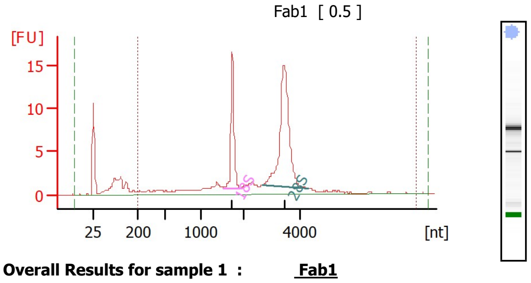


**Overall Results for sample 1 : Fab1**

| | |
|---|---|
| RNA Area: | 125.2 |
| RNA Concentration: | 85 ng/µl |
| rRNA Ratio [28s / 18s]: | 2.5 |
| RNA Integrity Number (RIN): | 9.2 (B.02.11) |
| Result Flagging Color: | |
| Result Flagging Label: | RIN: 9.20 |
| Corr. Area 1: | 106.1 |

**Fragment table for sample 1 : Fab1**

| Name | Start Size [nt] | End Size [nt] | Area | % of total Area |
|---|---|---|---|---|
| 18S | 1,524 | 1,915 | 17.7 | 14.1 |
| 28S | 2,673 | 4,266 | 43.7 | 34.9 |

**Region table for sample 1 : Fab1**

| Name | From [nt] | To [nt] | Corr. Area | % of Total | Color |
|---|---|---|---|---|---|
| DV200 | 200 | 8,000 | 106.1 | 85 | |

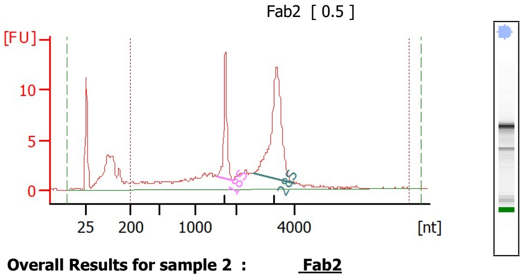


**Overall Results for sample 2 : Fab2**

| | |
|---|---|
| RNA Area: | 156.4 |
| RNA Concentration: | 106 ng/µl |
| rRNA Ratio [28s / 18s]: | 2.6 |
| RNA Integrity Number (RIN): | 8.2 (B.02.11) |
| Result Flagging Color: | |
| Result Flagging Label: | RIN: 8.20 |
| Corr. Area 1: | 121.0 |

**Fragment table for sample 2 : Fab2**

| Name | Start Size [nt] | End Size [nt] | Area | % of total Area |
|---|---|---|---|---|
| 18S | 1,525 | 1,882 | 14.8 | 9.5 |
| 28S | 2,597 | 4,013 | 38.6 | 24.7 |

**Region table for sample 2 : Fab2**

| Name | From [nt] | To [nt] | Corr. Area | % of Total | Color |
|---|---|---|---|---|---|
| DV200 | 200 | 8,000 | 121.0 | 77 | |

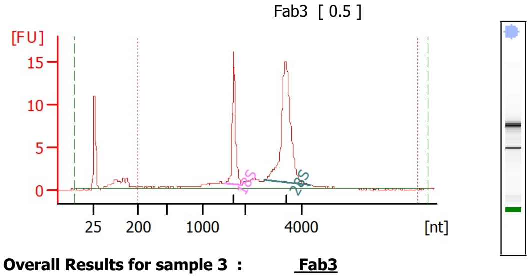


**Overall Results for sample 3 : Fab3**

| | |
|---|---|
| RNA Area: | 105.9 |
| RNA Concentration: | 72 ng/µl |
| rRNA Ratio [28s / 18s]: | 2.5 |
| RNA Integrity Number (RIN): | 9.4 (B.02.11) |
| Result Flagging Color: | |
| Result Flagging Label: | RIN: 9.40 |
| Corr. Area 1: | 94.0 |

**Fragment table for sample 3 : Fab3**

| Name | Start Size [nt] | End Size [nt] | Area | % of total Area |
|---|---|---|---|---|
| 18S | 1,544 | 1,934 | 18.0 | 17.0 |
| 28S | 2,684 | 4,294 | 44.7 | 42.2 |

**Region table for sample 3 : Fab3**

| Name | From [nt] | To [nt] | Corr. Area | % of Total | Color |
|---|---|---|---|---|---|
| DV200 | 200 | 8,000 | 94.0 | 89 | |

**S2 Fig. Confirmation of RNA quantity and purity for the FABP3-treated samples (Fab1, Fab2, and Fab3).** Sample Fab1 has a RIN of 9.2 and a 28S/18S ratio of 2.5. Sample Fab2 shows a RIN of 8.2 and a 28S/18S ratio of 2.6. Sample Fab3 displays a RIN of 9.4 and a 28S/18S ratio of 3.4. Standardized 28S/18S ratio, which is the relative abundance of the two ribosomal RNA subunits of around 2.0 indicates qualified RNA purity and integrity.

**Supplementary Figure 3 (S3 Fig)**

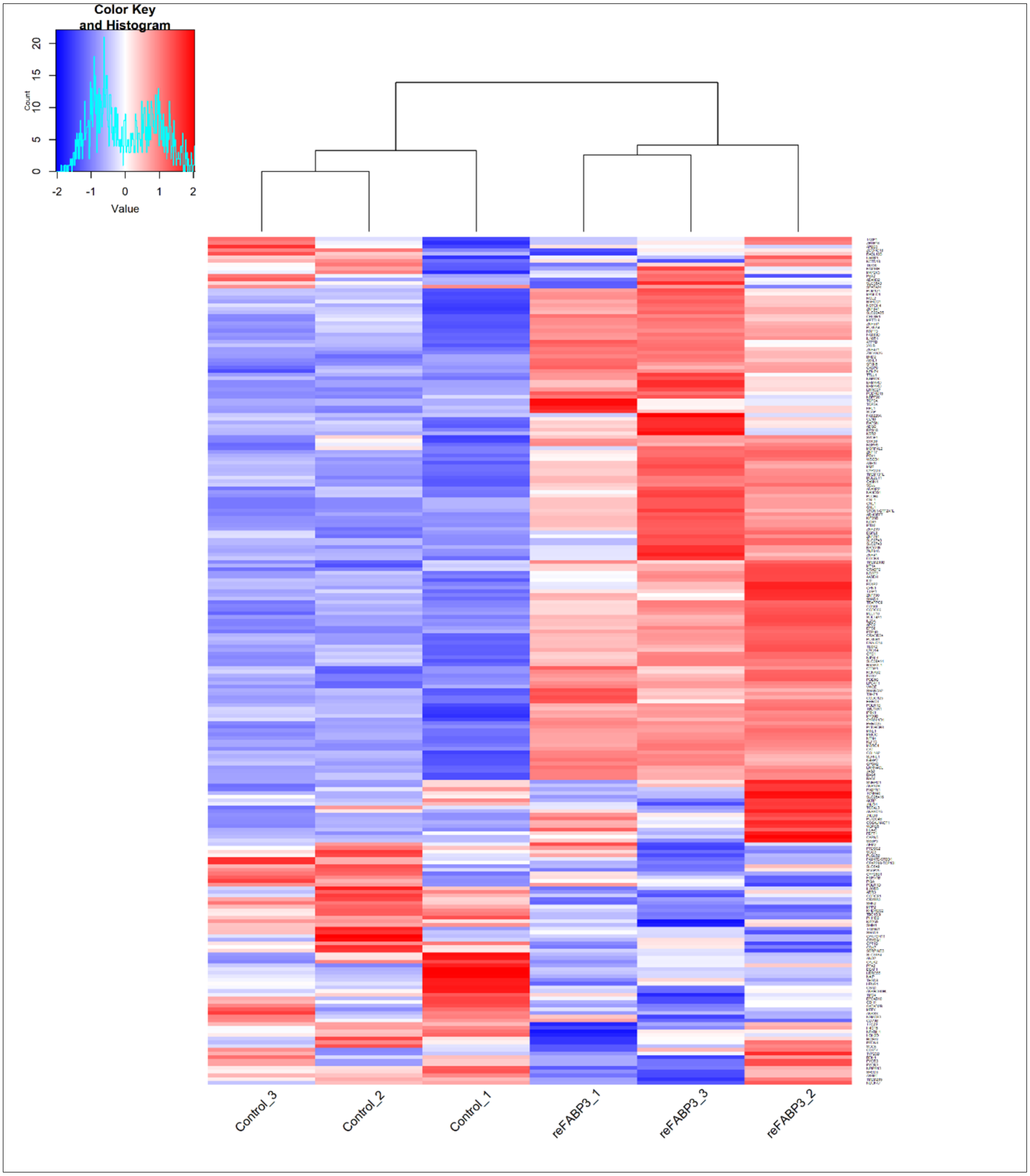


**S3 Fig. Gene expression heatmap showing clusters of samples from HUVECs treated with rhFABP3 (50ng/ml) for 6 hours vs. Vehicle groups**. The analysis was conducted on 231 protein-coding genes out of 15688 RNA seq genes tested for differential gene expression (DGE), with unadjusted p-values less than 0.05 and without the log (2) fold-change filtering criteria. N = 3. HUVEC = Human Umbilical Vein Endothelial Cells; rhFABP3 = recombinant human FABP3.

Supplementary Table 1 (S1 Table)

**S1 Table: RNA quantity and purity were assessed with the NanoDrop ND-1000**

| Sample Name | Nucleic Acid(ng/uL) | A260/A280 Ratio |
|---|---|---|
| Vehicle -1 | 302.076 | 1.872 |
| Vehicle-2 | 286.097 | 1.95 |
| Vehicle-3 | 248.249 | 1.928 |
| rhFABP3-1 | 278.835 | 1.985 |
| rhFABP3-2 | 300.089 | 1.93 |
| rhFABP3-3 | 207.784 | 1.922 |

**S1 Table. RNA quantity and purity were assessed with the NanoDrop ND-1000.**
